# A haptic illusion created by gravity

**DOI:** 10.1101/2023.01.19.524523

**Authors:** Laurent Opsomer, Benoit Delhaye, Vincent Théate, Jean-Louis Thonnard, Philippe Lefèvre

## Abstract

Haptic force estimation is a critical aspect of human dexterity. To manipulate objects or interact with a haptic interface successfully, the normal and tangential components of the contact forces produced by our fingertips must be carefully coordinated. The tight coupling between tangential and normal contact forces observed during voluntary movements indicates that the nervous system can estimate these forces with high accuracy. Here, we examined the influence of gravity on manual force production in an isometric task. We trained participants to produce isometric tangential forces with the thumb and index finger on a dynamometer held in precision grip. They were tasked with reproducing these forces in a normal gravitational environment (1 *g*) and during parabolic maneuvers creating phases of micro- (0 *g*) and hypergravity (1.6 *g*). The isometric task results showed that arm weight biases estimation of the forces exerted by the fingertips. Reproduced tangential forces were consistently larger downward than upward in 1 *g*. This asymmetry was reduced under microgravity and increased under hypergravity. Critically, the normal forces did not reflect this asymmetry, demonstrating that tangential forces were misestimated and pointing to a haptic illusion created by gravity. This gravitational effect on haptic force estimation may have implications for the design of prostheses incorporating force feedback, haptic devices for teleoperations, or haptic supports aimed at improving sensorimotor performance in space.

## INTRODUCTION

Fine manual force control is a fundamental aspect of dexterity, enabling humans to accomplish complex and eloquent tasks, from delicate surgeries to the teleoperation of a gigantic robotic arm while floating on board the International Space Station. Achieving such high-precision dexterity requires one to estimate the forces applied by the fingers to the tools they manipulate. Questions remain regarding how the nervous system combines tactile, proprioceptive, and internal efferent feedback to estimate these forces. Understanding the contribution of each sensorimotor modality to force estimation is of primary importance for the design of prostheses that will incorporate sensory feedback and of haptic supports for remote operations including telesurgery.

During object manipulation, grasping forces must be finely controlled to prevent accidental slips while minimizing muscle fatigue. Slips occur when the tangential component of the contact force (a friction force) exceeds some threshold proportional to the normal component of the contact force (Coulomb’s law). When holding an object in precision grip (i.e. between the thumb and the index finger), one can ensure that this threshold is not exceeded by tuning the grip force (GF), which is applied perpendicularly to the contact surfaces, to the load force (LF), which is produced by the tangential forces applied by each finger^1,2^. Many studies have examined the coordination between LF and GF during object manipulation to probe human dexterity. In a large variety of tasks (arm movements^1,3,4^, locomotion^5^, controlled collisions^6–8^, etc.), tight coupling between LF and GF is observed, suggesting that the nervous system estimates LF online with good accuracy during voluntary movements, which allows precise anticipative and reactive adjustments of GF.

Despite decades of studies in the field, the exact contribution of each sensorimotor modality to LF estimation remains unclear. Notwithstanding, it is clear that tactile feedback transmitted by cutaneous mechanoreceptors plays a primordial role^1,9–13^. These receptors are sensitive to skin deformation and partial slips, and their input allows precise estimation of tangential and normal contact forces^14–18^. Recent studies have shown that stretching the skin artificially during object manipulation can alter the perception of LF and thus the control of GF^19,20^. The nervous system can also estimate LFs based on the activities of muscles involved in the production of these forces. Such muscular activity can be sensed via proprioceptive receptors (muscle spindles and tendon organs)^21–23^ or predicted based on motor commands sent to the muscles^24,25^. Here, we examined the contribution of muscle activity to LF estimation by manipulating the contribution of gravity to LF production.

To obtain accurate estimates of manual forces based on muscle activity, the nervous system must take gravity into account. In particular, limb weight modulates the muscular effort required to produce a given output force. For example, applying a downward force on a haptic interface requires less effort than applying the same force upward, due to gravity. Previous studies have reported that scaling between GF and LF is not affected by loading of the arm with ballasts or elastic straps during rhythmic arm movements^26,27^. Furthermore, tight GF-LF coupling can be maintained across the various gravito-inertial levels experienced during parabolic flight after a short adaptation period ^26,28,29^. Thus, the nervous system can account for muscular effort when estimating LF *during active arm movements.* However, because the sensory feedback information available for force estimation during isometric task performance differs fundamentally from that available during movement, it is unknown whether force estimation is also accurate *in the absence of movement*, that is, during isometric force production.

During voluntary movements, proprioceptive and visual feedback about arm kinematics can be combined to infer manipulation forces^30,31^. In unfamiliar environments, feedback about movement errors helps correct motor commands and update internal representations of limb dynamics ^32,33^. In microgravity, for instance, motor commands adjust rapidly to reproduce movements learned in normal gravity ^34–37^. The new internal representation of limb dynamics can then be used to update the feedforward mechanisms participating in LF estimation ^38–40^.

In isometric conditions, motion feedback and movement errors are not available to aide in LF estimation or to update internal models. In this context, the nervous system can rely only on tactile and proprioceptive force feedback and on “sense of effort” based on efference copies ^22,24,41,42^. Therefore, force estimation may be less accurate and more affected by unfamiliar gravito-inertial environments when contact forces are produced isometrically compared with force estimation during active movements. To test this hypothesis, we studied the ability of human participants to reproduce memorized isometric load forces during exposure to various gravito-inertial levels (G-levels) induced by parabolic flight.

Participants were trained on the ground to produce upward and downward isometric LFs of a specific magnitude on a static dynamometer held in precision grip. They were asked to reproduce these forces without explicit feedback during parabolic maneuvers producing distinct G-level phases, which constituted microgravity (0 *g*) and hypergravity (1.6 *g*) conditions. We found that gravity was associated with LF production bias such that the magnitude of LF reproduced was consistently larger in the downward direction than in the upward direction in normal gravity. Importantly, this bias was reduced in microgravity and more pronounced in hypergravity. In contrast, GF values were similar in both LF directions regardless of G-levels, providing evidence that LF was not estimated accurately by the nervous system when adjusting GF as a function of LF to secure the grasp. The results of this study thus suggest that gravity biases force estimation during isometric force production, creating a haptic illusion.

## RESULTS

### Establishment of the experimental model

Participants (N = 11) were trained to apply upward and downward isometric LFs of around 5 N on a dynamometer fixed to a support (see Methods). They held a dynamometer in a precision grip (Figure 1A) and applied an isometric LF by pulling the dynamometer upward or downward. At regular intervals, the target LF was communicated via an auditory feedback cue, namely a beep that was emitted whenever the LF magnitude was within the range of 4–6 N. LF was computed as the absolute value of the resultant force applied by both fingers along the vertical axis, tangentially to the contact surfaces. Applied GF was computed as the mean of the normal forces applied by each finger.

**Figure 1.**
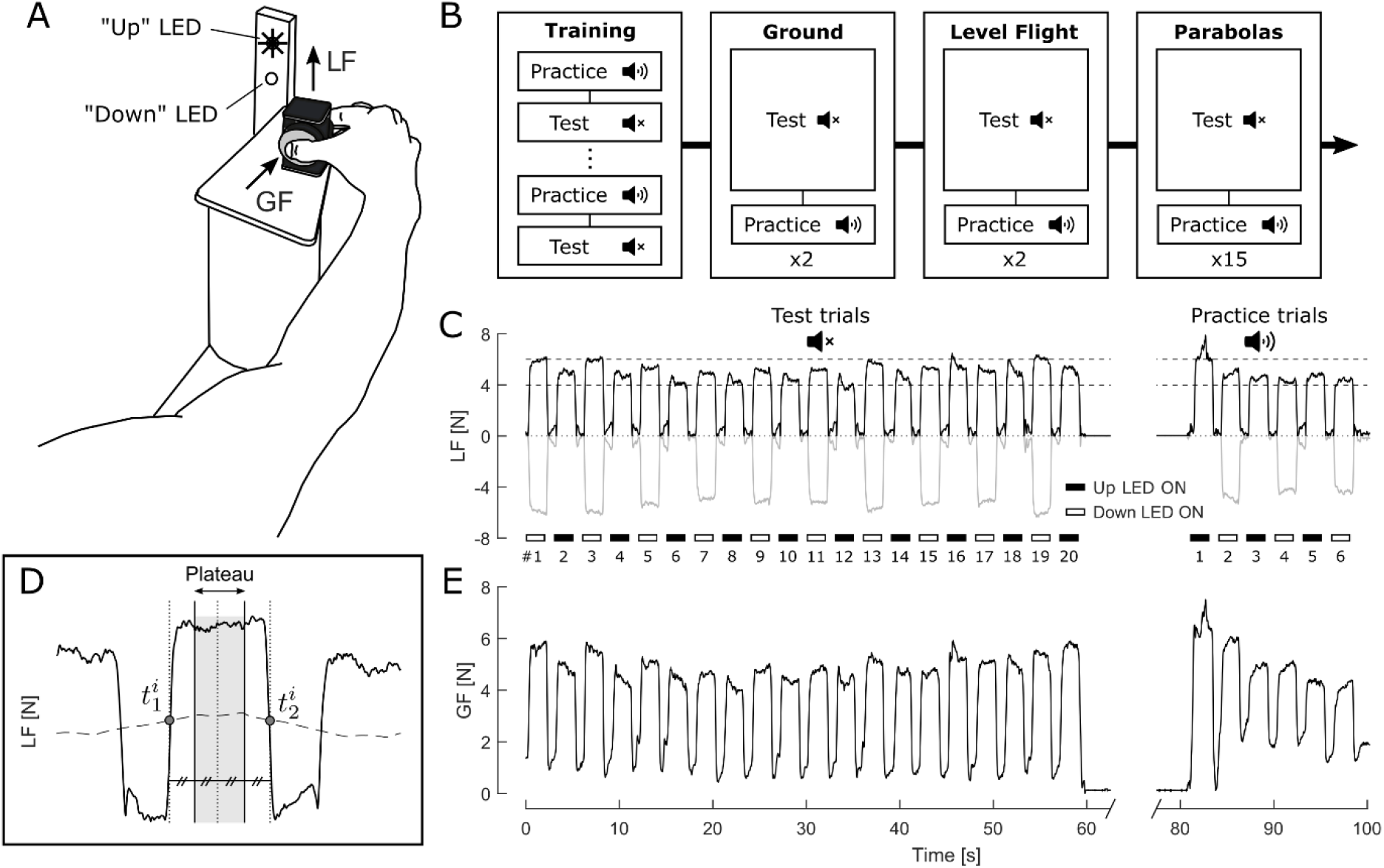
Experimental design and data analysis. (A) Task: Participants held a static dynamometer between their thumb and index finger and were instructed to produce upward and downward isometric load forces (LFs) of 5 ± 1 N. An LED display indicated the direction of LF to produce. (B) Protocol: Participants were first trained on the ground to produce the desired LF. During practice trials, a beep was emitted when the produced LF was within the target bounds ([4 N, 6 N]). During test trials, no auditory feedback was given. They were asked to reproduce the memorized forces on the ground, during level flight, and during parabolic maneuvers producing phases of micro- and hypergravity. (C) LF during a typical sequence of test and practice trials performed on the ground. The gray line shows the signed values of LF (negative for downward forces). The dashed horizontal lines indicate the target LF (5 ± 1 N). (D) Magnified view of LF during one trial from panel C. Individual trials were delimited by the points 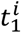 and 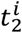 defined as the intersections between LF (plain line) and a down-scaled moving average of LF computed with a 3-s sliding window (dashed line). For each trial, the plateau phase was defined as a window centered between 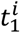 and 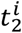 (shaded gray area). (E) Grip force (GF) measured in the trials shown in panel C.

Trained participants reproduced the learned LF, first on the ground (≤24 h before the flight) and then onboard the aircraft during level flight in a steady 1-*g* environment, before undertaking the experiment during parabolic maneuvers (Figure 1B). On the ground and during level flight, they performed two sequences of 20 trials without auditory feedback (test trials, Figure 1C), alternating between downward and upward LFs. They performed 6 additional trials with auditory feedback after each such sequence to remind them of the target (practice trials, Figure 1C). Each trial lasted 2 s with a 1-s inter-trial break. The force data were averaged within the plateau phase of each trial (Figure 1D; see Methods).

### Isometric LF reproduction in normal gravity

We focus here on data obtained during level flight (Figure 2), which were similar to the ground data (see Figure S1). During the test trials, LF remained on average within the target bounds yielding a mean LF [± standard deviation (SD)] of 4.98 ± 1.15 N (Figure 2A). LF decreased gradually across the 20 trials of each sequence, by about 18% on average. A concomitant decrease in GF of similar magnitude was observed (Figure 2B). We observed a strong temporal correlation between GF and LF for all test sequences (r = 0.93 ± 0.05, mean ± SD) consistent with the GF-LF coupling typically observed in object manipulation tasks.

**Figure 2.**
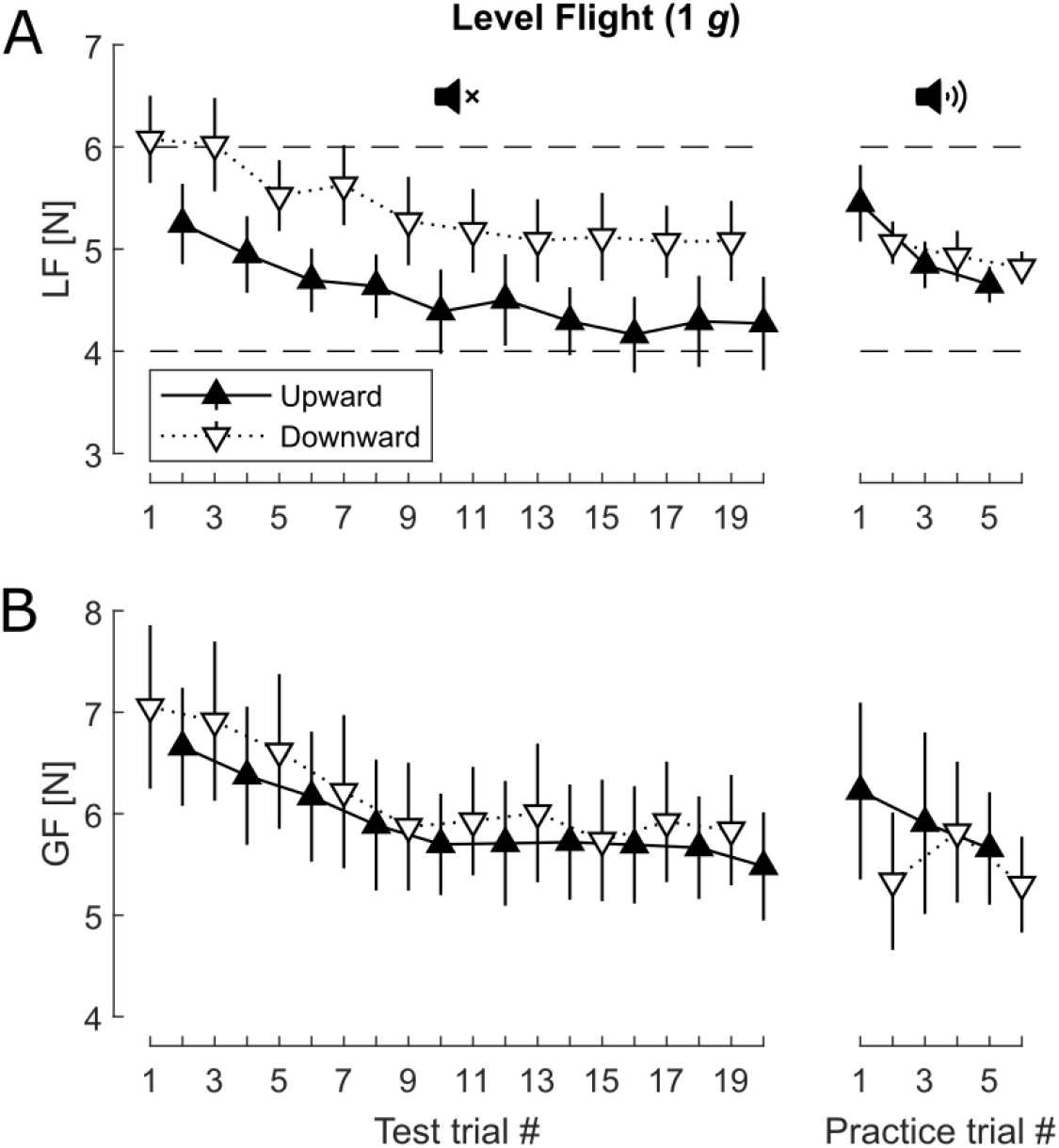
Average force data during level flight. Evolution of LF (A) and GF (B) across test and practice trials performed during level flight. The two sequences were averaged by participant and trial number. The figure shows the means (upward/downward pointing triangles for upward/downward LF) ± 1 standard error of the mean (SEM; error bars) across participants (N = 11). The horizontal dashed lines in panel A show the LF target zone.

Strikingly, up/down asymmetry in LF was observed. LF was significantly larger in the downward direction than in the upward direction, by 19% (0.86 N) on average (t_10_ = −3.56; *p* < 0.01). Despite tight temporal coupling between LF and GF, GF remained similar for the two LF directions (Figure 2B; t_10_ = −1.06; *p* = 0.31). During the practice trials performed with auditory feedback, both LF (t_10_ = −0.74; *p* = 0.48) and GF (t _10_ = 0.94; *p* = 0.37) were similar in the two directions, as expected.

### G-level produces distinct effects on LF asymmetry versus GF asymmetry

When the participants repeated the same task during parabolic maneuvers, during which the gravito-inertial level (G-level) varied from 0 *g* to around 1.6 *g* (Figure 3A), they reproduced the LF that had been learned in 1 *g* in both the microgravity (0 *g*) and hypergravity (1.6 *g*) conditions with good accuracy (Figure 3B), even though auditory feedback was not given during the parabolas. Nevertheless, a clear effect of G-level on up/down LF asymmetry was observed. During microgravity phases (trials 1–6), when G-level (and thus arm weight) was close to zero, LF asymmetry was reduced compared with LFs produced in 1 *g* during level flight. Subsequently, as G-level increased from 0 to 1.6 *g* (trials 7–12), the difference between downward and upward LFs increased substantially, exceeding a 20% differential on average. This asymmetry decreased progressively as the G-level returned to 1 *g* (trials 12–20). This pattern was observed from the first to the last parabola (see Figure S2 and Methods) and was observed for *experienced* participants (N = 6, see Methods) as well as for *inexperienced* participants (N = 5) who were being exposed to parabolic maneuvers for the first time (Figure S3).

**Figure 3.**
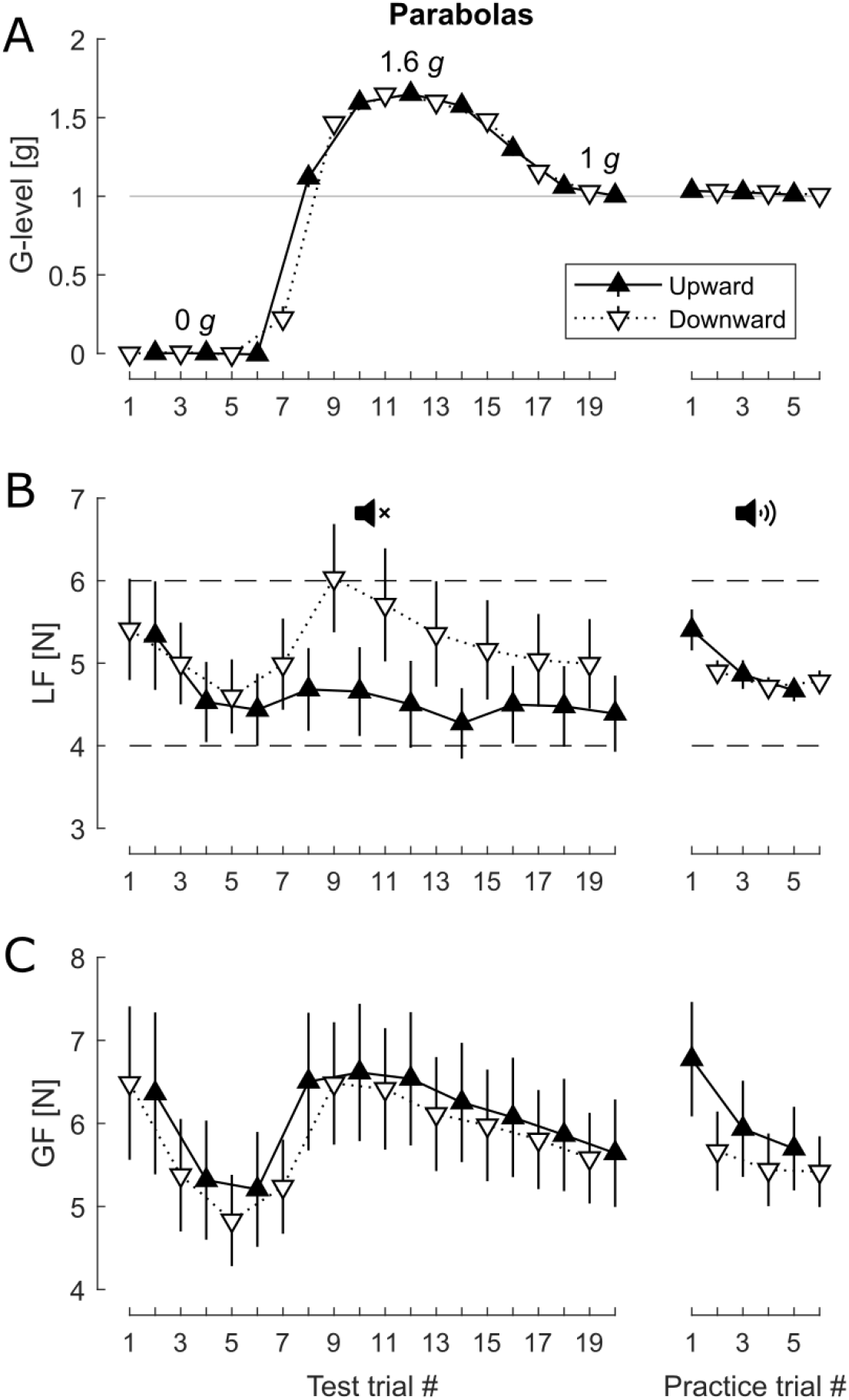
Average force data during the parabolas. Evolution of G-level (A), LF (B) and GF (C) across test and practice trials. The data obtained in 15 parabolas performed by each participant were averaged by participant and trial number. The figure shows the means (upward/downward pointing triangles for upward/downward LF) ± SEM (error bars) across participants (N = 11). The horizontal dashed lines in panel B show the LF target zone.

In contrast to LFs, GFs remained symmetrical at all G-levels, being similar for upward LFs and downward LFs (Figure 3C), resulting in a decoupling between LF and GF magnitudes. GF appeared to follow LF inter-trial variations well for downward trials, but less so for upward trials. Notwithstanding, a strong temporal correlation was maintained between GF and LF and the mean correlation coefficient (±SD) during the parabolas was 0.90 ± 0.05. This coefficient was stable across parabolas. Similar to the behavior observed during level flight, participants adjusted GF as a function of LF, but they failed to account for the directional bias in LF production modulated by G-level.

To quantify the effects of G-level and direction on LF and GF, we divided the test trials into three distinct bins corresponding to 0-, 1-, and 1.6-*g* gravito-inertial accelerations (see Methods). The mean LF, GF, and GF/LF ratio values obtained are shown in Figure 4 (upper panels) together with the mean differences in values between upward and downward trials (lower panels). LF was globally significantly greater in the downward direction than in the upward direction (F_1,10_ = 26.5, *p* < 0.001; Figure 4A); this asymmetry was modulated by G-level, as indicated by a significant interaction effect between G-level and direction (F_2,20_ = 5.34, *p* = 0.01; Fig. 4B). There was no main effect of G-level on LF (F_2,20_ = 0.59, *P* = 0.57). Conversely, GF did not differ significantly between upward and downward trials (Figure 4C-D; F_1,10_ = 0.03, *p* = 0.87). GF tended to increase with G-level, but the effect was not significant (F_2,20_ = 2.54, *p* = 0.10) and there was no significant interaction between G-level and direction for GF (F_2,20_ = 1.33; *p* = 0.29). Due to decoupling between LF and GF magnitudes, the GF/LF ratio was not constant across directions and G-levels (Figure 4E). Rather, we observed significant main effects of both G-level (F_2,20_ = 9.33, *p* < 0.005) and direction (F_1,10_ = 10.2; *p* < 0.01) on the GF/LF ratio, as well as significant interaction between G-level and direction for GF/LF ratio (F_2,20_ = 7.84, *p* < 0.005; Figure 4F). Scaling between GF and LF was thus gravity-dependent.

**Figure 4.**
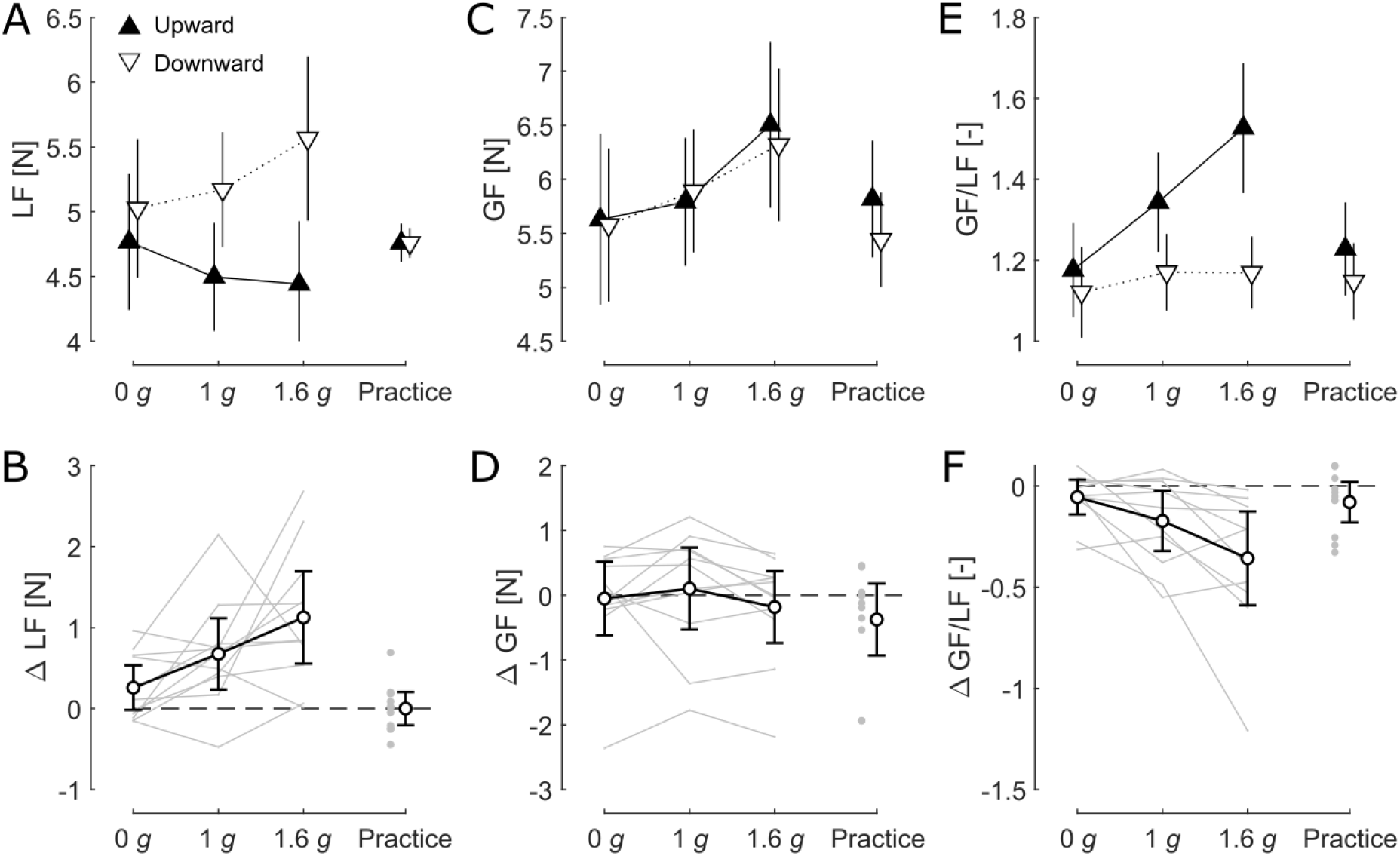
Up/down asymmetry as a function of G-level. (A, C, E) Mean LF, GF, and GF/LF values in each G-level bin during practice and test trials. Triangle markers and error bars show the mean ± 1 SEM across participants (N = 11). (B, D, F) Mean differences between downward and upward trials for LF, GF, and GF/LF ratio. Error bars with caps show the 95% confidence intervals. Gray lines show data from each participant. The 1-g and practice bins include the level-flight data shown in Figure 2.

## DISCUSSION

The present study shows that gravity biases isometric force production, as reflected by a gravity-dependent up-down asymmetry of reproduced LF. Unexpectedly, when participants were instructed to reproduce a vertical LF that had been practiced on the ground (1 *g*) with symmetry relative to zero (±5 N), they failed to reproduce the symmetry practiced in training, even in the exact same gravity condition. Reproduced LFs had consistently larger downward magnitudes than upward magnitudes in a 1 *g* condition. This asymmetry increased in hypergravity and disappeared in microgravity during parabolic maneuvers, and thus we deduce that it was directly related to arm weight. Critically, GF, which is supposed to be proportional to LF to prevent accidental slips^1^, had always similar magnitudes whether LF was produced upward or downward. In other words, GF was not accurately tuned to LF, indicating that LFs were not estimated accurately by the nervous system. Hence, we argue that our data reflect a haptic illusion—an overestimation of upward forces and/or an underestimation of downward forces—that is caused by and modulated by gravity.

The present results are compatible with the hypothesis that the nervous system estimates manual forces by combining tactile feedback with a sense of effort related to muscle activity^24,43,44^. By detecting skin deformations and partial slips, tactile feedback alone should provide accurate tangential force feedback unbiased by gravity^14–17^, but tactile feedback is also subject to uncertainties. For instance, a given tangential force can in general lead to distinct patterns or amplitudes of skin deformations (and thus afferent responses) depending on the normal component of the contact force as well as on the frictional properties of the contact interface ^15,16,45^. Combining tactile feedback with a sense of effort should reduce these uncertainties to some degree. However, muscular effort is necessarily biased by gravity in that producing a 5-N LF in opposition to gravity (upward) requires more effort than producing the same LF in the direction of gravity (downward). If muscular effort contributes to force estimation, upward LFs would be expected to be overestimated relative to downward LFs. Furthermore, background G-level modulates the tonic muscle activity required to compensate for limb weight. Underestimating this tonic activity in hypergravity would lead to an under-compensation of arm weight, which would increase the asymmetry between upward and downward LFs, as observed in our data.

Muscular effort can be sensed via internal feedback about motor commands or via proprioceptive feedback conveyed by muscle spindles and tendon organs. Both motor commands^24,41^ and proprioception^21–23^ have been shown to influence the conscious perception of forces. Anisotropies in haptic force perception have been observed in the horizontal plane previously, with manual forces produced in directions requiring larger effort (due to larger limb impedance) being perceived as larger^43,44^. The present data reveal an anisotropy in haptic force estimation in the vertical plane (up/down asymmetry) caused by gravity that affects GF-LF coupling.

The present findings indicating that a sense of effort may influence grip control in isometric conditions contrasts with the results of several past studies that have examined coupling between GF and LF during active arm movements. In those studies, the authors observed a constant scaling between GF and LF during rhythmic arm movements in altered gravity^26,29^ or when the arm was loaded with a ballast^26^ or an elastic band ^27^, indicating that GF control can be tuned accurately to LF independently of the muscle activity generated to move the arm. This dissimilarity relative to our results suggests that the additional visual and kinesthetic feedback available during unconstrained arm movements may improve LF estimation significantly. Internal inverse models could be used to infer manipulation forces from the kinematics of the manipulated object ^30,31^. In agreement with this hypothesis, altered visual feedback of object motion has been shown to affect the GF-LF coupling ^46–48^. In addition, movement errors can be used to enable rapid adaptation of motor commands to the novel dynamics of an unfamiliar environment. In parabolic flight, a few parabolas are typically required to adapt movement kinematics ^37,49^ and to stabilize GF-LF coupling ^28,29,50^. The absence of movement errors might be one factor explaining the absence of significant learning in our paradigm.

Degradation in the ability to reproduce isometric forces in micro-^51^ and hypergravity has been described previously ^52,53^. In those studies, consistent overshooting of the remembered target force was observed in altered gravity in both the horizontal and vertical planes. The absence of such direction-independent overshooting during the parabolas in the present study could be explained by the presence of higher precision tactile feedback wherein a precision grip was employed rather than a power grip as had been used in the previous studies. A precision grip allows for finer cutaneous feedback from the dense population of mechanoreceptors in the fingerpads and from the joint, tendon, and muscle receptors of the fingers. Additionally, the task performed was more complex in the aforementioned studies, with participants being expected to memorize 3 to 5 different target magnitudes presented in 4 to 8 different directions.

The present work opens the door to many future experiments that should aim to clarify the exact mechanisms underlying the presently demonstrated gravity-dependent haptic illusion. Future work could test the respective contributions of shoulder, elbow, wrist, and finger muscles to LF estimation, with the use of ballasts, elastic chords, and counterweights attached to various points of the arm during isometric force production. Such experiments can allow the effects of muscle activity to be distinguished from other effects specific to the conditions of microgravity and hypergravity, conditions that also affect vestibular inputs, postural control ^35,36^, spatial orientation ^54–56^, limb position sense ^57–59^, and stress ^60^. Our protocol did not allow us to probe the participants’ conscious perception of forces. Studying how gravity influences force perception, in addition to the estimations performed by the sensorimotor system in the context of GF control, could improve our understanding of the phenomenon. Although GF control has been shown to be unaffected by some perceptual illusions, such as the size-weight^61,62^ or material-weight^63^ illusions, we showed here that GF control can be perturbed by gravity-dependent haptic illusion. In the same vein, GF control can be affected by illusions of increased stiffness caused by artificial skin stretch^20^.

In conclusion, our study indicates that gravity biases force estimation during isometric object manipulation and, consequently, that force estimation can be altered in unfamiliar gravity conditions. This gravity-related haptic illusion could be considered when designing haptic devices used to support telesurgeries^64^ or other remote operations, or haptic supports aimed at improving sensorimotor performance in astronauts^65^. One could implement direction-dependent gains, such that the force output would reflect more accurately the intention of the user, rather than following the actual force produced. For haptic devices to be used in aircraft, in rockets, or on board the International Space Station, these gains could be adjusted as a function of the gravito-inertial level given that our results indicate that gains optimized for 1-*g* conditions on the Earth’s surface may not be optimal in reduced or increased gravity settings. Gravity-dependent gains could help minimize errors in force production when visual feedback about performance is unavailable or unreliable—errors that can have dramatic consequences in these contexts.

## Acknowledgments

The authors would like to thank the participants of this study. This work was supported by a grant from the European Space Agency (Prodex) and the Belgian Federal Government. The manuscript was edited by a professional scientific editor at Write Science Right.

## Author contributions

V.T., B.D., J-L.T and P.L. designed and conducted the experiment. L.O. and V.T. analyzed the data. L.O. wrote the paper and prepared the figures. All authors helped revise and edit the paper.

## MATERIALS AND METHODS

### Subjects

A group of 12 healthy adult volunteers (age, 23—58 years; 9 males, 3 females) participated in this experiment; 6 participants had participated in at least one parabolic flight prior to this study, while the other 6 participants had never experienced parabolic flights before. Data from one participant was excluded (explanation below in the Experimental procedure section). All participants gave their informed consent and received approval for parabolic flight in a National Center for Aerospace Medicine class II medical examination. The experiment complied with the European Space Agency ethical and biomedical requirements for experimentation on human subjects (ESA Medical Board Committee) and was approved by the local French ethics committee (CPP) in charge of reviewing life science protocols in accordance with French law.

### Experimental model

The experiment was performed during the 58^th^ ESA parabolic flight campaign on board the A-300 zero-*g* aircraft. Each parabolic maneuver began with 20-s of hypergravity (pull-up phase) followed by 22-s of microgravity (0 *g*), before another 20-s period of hypergravity (pull-out phase; 1.6 *g*) ensued and then there was a progressive transition back to a 1-*g* gravito-inertial level. The beginning and end of each microgravity phase were announced by the pilot as the “injection” and “pull-out” phases, respectively. One flight was performed each day for three consecutive days; a sequence of 31 parabolas was performed in each flight.

#### Task

The experimental task consisted of applying the LF, an isometric tangential vertical force of a specified magnitude (5 ± 1 N) and direction (upward or downward), on a static dynamometer held in precision grip. Participants were instructed to pull the dynamometer either upward or downward, vertically with respect to the aircraft reference frame (and therefore always aligned with the gravito-inertial force). The force had to be maintained for 2 s.

#### Training

Between 1 and 14 days prior to flight, the participants were trained to memorize and reproduce the desired upward and downward LFs on the dynamometer. The training consisted of sequences of *practice* and *test* trials. During practice trials, auditory force feedback was given as a beep that emitted whenever the LF magnitude was within the target bounds (4–6 N). During test trials, the participants were asked to reproduce the memorized upward or downward LF, and no feedback was given. Each participant performed at least four training sequences of 60 trials divided into blocks of 6 practice trials followed by 6 test trials, always alternating between an upward trial and a downward trial. Each trial lasted 2 s (time interval between the onset and offset of the LED target) and the time interval between successive trials was 1 s. A short break was imposed every 30 trials.

#### Experimental procedure

Once properly trained, participants performed the actual experiment, segmented into sequences of 26 trials. Each sequence started with 20 test trials and ended with 6 practice trials after a 20-s break to maintain good memorization of the forces to be reproduced. Trials always alternated between an upward and a downward force, as illustrated in Figure 1B. As during training, every trial lasted 2 s, with an inter-trial interval of 1 s. The participants performed at least two of these sequences on the ground, then performed two additional sequences on board the aircraft during level flight (1 *g*). Finally, they performed identical sequences during 15 consecutive parabolas. During the parabolas, the sequence started at the injection point (onset of 0-*g* phase) and ended in 1-g after the pull-out phase. On average, participants performed 6 test trials in 0 *g*, ~5 test trials at >1.5 *g*, as well as ~3 test trials in 1 *g* per parabola. The other test trials were performed during the transition phases of the parabolas. All practice trials were performed in 1 *g*.

We rejected the trials that were performed in the wrong direction and those for which the standard deviation of the LF exceeded 20% of the mean during the plateau phase (<1% of trials). Due to technical issues, data could not be acquired during three parabolas for four participants and during five parabolas for two other participants. Furthermore, data could not be acquired during one levelflight block for two participants. In addition, the auditory feedback did not work for one participant on board the aircraft during the practice trials; this participant was removed from the data set.

### Data collection

Duplicate experimental setups allowed data to be acquired from two participants simultaneously during each flight. Because each participant performed the experiment during 15 consecutive parabolas, four participants were tested per flight. The participants were seated and restrained in pairs of side-by-side chairs. In front of each participant, a dynamometer was fixed on a horizontal platform sitting vertically with respect to the aircraft floor. Each dynamometer (GLM, Arsalis, Louvain-La-Neuve, Belgium) was equipped with two force-torque sensors (40 mm diameter, Mini 40 F/T transducers, ATI Industrial Automation, Apex, NC, USA), one for the thumb and one for the index finger. The final distance (50 cm) and orientation of the dynamometer relative to the chair were chosen such that all participants were comfortable when grasping the device in precision grip (Figure 1A). The arm used for grasping (right arm) was not supported during the experiment. Two LEDs indicated the direction of the LF to be applied on the dynamometer (upward or downward).

On board the aircraft, G-level was measured with a three-dimensional accelerometer (ADXL330, Analog Devices). Force and acceleration signals were all sampled at 1000 Hz and synchronized thanks to a signal conditioner (Arsalis, Louvain-La-Neuve, Belgium) and a data acquisition system (National Instruments, USA). Offsets of force signals were cancelled before each test sequence so that they were at zero when no force (other than gravity) acted on the sensors. LF was computed as the absolute value of the sum of the vertical component of the tangential forces applied by each finger. GF was computed as the mean of the normal component of the forces applied by each finger.

#### Data analysis

The data were processed in custom software written in Matlab (Mathworks, USA). Because force offsets were cancelled in a 1 *g* setting (see previous paragraph), force signals obtained during parabolas were corrected to avoid a bias induced by the weight of the sensor plate varying as a function of G-level. This correction consisted in subtracting a G-level-dependent offset, *C* = *m_s_*(1 – *G*) · 9.81, from the vertical component of the force measured by each sensor. *m_s_* is the mass of the sensor plate (0.023 kg) and *G* is the gravito-inertial level. Force and accelerometer data were digitally low-pass filtered with a zero phase-lag Butterworth filter of order four with cut-off frequencies of 40 Hz and 5 Hz, respectively.

Because participants were instructed to apply a vertical force and auditory feedback was related to that vertical component during practice trials, we did not consider the horizontal component of tangential forces applied to the dynamometer. The vertical component of the tangential force constituted the LF and accounted for, on average, 98.0 ± 2.6% of the total tangential force.

LF was used to segment the data into individual trials, as illustrated in Figure 1D. A moving-average of LF was computed using a 3-s sliding window and then down-scaled by a factor of 0.8. This down-scaling was performed such that the moving-average crossed the LF signal at around half the value reached by LF during the plateau phase of each trial. The crossing points of this down-scaled moving-average with the LF signal, 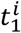 and 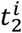, were used to define the plateau phase in each trial: the plateau phase was defined as a time window located halfway between 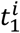 and 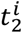 and having a width of 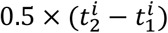. On average, the duration of the resulting plateau phases was 971 ± 123 ms (mean ± SD). Mean LF and GF values were computed within each plateau phase.

### Statistical analyses

Statistical analyses were performed in R using the *ez* package. For analyses of ground and level-flight data, two-sided paired t-tests were used to compare upward and downward test trials. We also used two-sided paired t-tests to compare upward and downward practice trials (the 1^st^ and 2^nd^ practice trials of each sequence were not included in this analysis to focus on stable performance with explicit auditory feedback). For analyses of the parabola data, the test trials were divided into three bins: a 0-*g* bin for trials performed at a mean G-level between −0.05 and 0.05 *g*; a 1-g bin for trials performed during level flight and at the end of each parabola at a mean G-level between 0.9 and 1.1 *g;* and a 1.6-*g* bin for trials performed at a mean G-level greater than 1.5 *g*. The trials performed during the transition phases of the parabolas were not included in these bins. We checked whether GF and LF changed across parabolas by comparing the first parabola to the last five parabolas in a 3-way repeated-measures ANOVA with factors of Parabola (first/last five), Direction (upward/downward) and G-level (0 *g*/1 *g*/1.6 *g*). During the first parabola, the participants tended to produce greater LFs (and GFs) compared with subsequent parabolas. Because the effect was marginal (F_1,10_ = 3.86, *p* = 0.07) and independent from G-level (F_2,20_ = 0.81, *p* = 0.41) and direction (F_1,10_ = 2.91, *p* = 0.08), all parabolas were pooled together. The effects of direction (upward/downward) and G-level (0 *g*/1 *g*/1.6 *g*) on LF and GF were then tested in 2-way repeated measures ANOVAs. The assumption of sphericity was verified with Mauchly’s test. When this assumption was violated (*p* < 0.05), *p* values were subjected to Greenhouse-Geisser correction.

To study GF-LF temporal coupling, we computed the correlation coefficient between GF and LF signals taken between the onset of the first test trial and the offset of the last test trial of each 20-trial sequence. The correlation coefficient was computed with the Matlab function *corrcoef*.

## References

1. Johansson, R.S., and Westling, G. (1984). Roles of glabrous skin receptors and sensorimotor memory in automatic control of precision grip when lifting rougher or more slippery objects. Exp. Brain Res. 56, 550–564. 10.1007/BF00237997.

2. Westling, G., and Johansson, R.S. (1984). Factors influencing the force control during precision grip. Exp. Brain Res. 53, 277–284. 10.1007/BF00238156.

3. Flanagan, J.R., and Wing, A.M. (1995). The stability of precision grip forces during cyclic arm movements with a hand-held load. Exp. Brain Res. 105, 455–464. 10.1007/BF00233045.

4. Flanagan, J.R., and Wing, A.M. (1993). Modulation of grip force with load force during point to point arm movements. Exp. Brain Res. 95, 131–143.

5. Gysin, P., Kaminski, T.R., and Gordon, A.M. (2003). Coordination of fingertip forces in object transport during locomotion. Exp. Brain Res. 149, 371–379. 10.1007/s00221-003-1380-1.

6. White, O., Thonnard, J.-L., Wing, A.M., Bracewell, R.M., Diedrichsen, J., and Lefèvre, P. (2011). Grip force regulates hand impedance to optimize object stability in high impact loads. Neuroscience 189, 269–276. 10.1016/j.neuroscience.2011.04.055.

7. Bleyenheuft, Y., Lefèvre, P., and Thonnard, J.-L. (2009). Predictive mechanisms control grip force after impact in self-triggered perturbations. J. Mot. Behav. 41, 411–417. 10.3200/35-08-084.

8. Delevoye-Turrell, Y.N., Li, F.X., and Wing, A.M. (2003). Efficiency of grip force adjustments for impulsive loading during imposed and actively produced collisions. Q. J. Exp. Psychol. Sect. A Hum. Exp. Psychol. 10.1080/02724980245000025.

9. O’Shea, H., and Redmond, S.J. (2021). A review of the neurobiomechanical processes underlying secure gripping in object manipulation. Neurosci. Biobehav. Rev. 10.1016/j.neubiorev.2021.01.007.

10. Witney, A.G., Wing, A.M., Thonnard, J.-L., and Smith, A.M. (2004). The cutaneous contribution to adaptive precision grip. Trends Neurosci. 27, 637–643. 10.1016/j.tins.2004.08.006.

11. Augurelle, A.-S., Smith, A.M., Lejeune, T., and Thonnard, J.-L. (2003). Importance of Cutaneous Feedback in Maintaining a Secure Grip During Manipulation of Hand-Held Objects. J. Neurophysiol. 89, 665–671. 10.1152/jn.00249.2002.

12. Nowak, D.A., Hermsdörfer, J., Glasauer, S., Philipp, J., Meyer, L., and Mai, N. (2002). The effects of digital anaesthesia on predictive grip force adjustments during vertical movements of a grasped object. Eur. J. Neurosci. 14, 756–762. 10.1046/j.0953-816X.2001.01697.x.

13. Okamoto, S., Wiertlewski, M., and Hayward, V. (2016). Anticipatory Vibrotactile Cueing Facilitates Grip Force Adjustment during Perturbative Loading. IEEE Trans. Haptics 9, 233–242. 10.1109/TOH.2016.2526613.

14. Paré, M., Carnahan, H., and Smith, A.M. (2002). Magnitude estimation of tangential force applied to the fingerpad. Exp. Brain Res. 142, 342–348. 10.1007/S00221-001-0939-Y.

15. Barrea, A., Delhaye, B.P., Lefèvre, P., and Thonnard, J.-L. (2018). Perception of partial slips under tangential loading of the fingertip. Sci. Rep. 8. 10.1038/s41598-018-25226-w.

16. Delhaye, B.P., Jarocka, E., Barrea, A., Thonnard, J.-L., Edin, B.B., and Lefèvre, P. (2021). High-resolution imaging of skin deformation shows that afferents from human fingertips signal slip onset. Elife 10. 10.7554/ELIFE.64679.

17. Delhaye, B.P., Lefèvre, P., and Thonnard, J. (2014). Dynamics of fingertip contact during the onset of tangential slip. J. R. Soc. Interface 11, 20140698. 10.1098/rsif.2014.0698.

18. Birznieks, I., Macefield, V.G., Westling, G., and Johansson, R.S. (2009). Slowly adapting mechanoreceptors in the borders of the human fingernail encode fingertip forces. J. Neurosci. 29, 9370–9379. 10.1523/JNEUROSCI.0143-09.2009.

19. Quek, Z.F., Schorr, S.B., Nisky, I., Okamura, A.M., and Provancher, W.R. (2014). Augmentation of stiffness perception with a 1-degree-of-freedom skin stretch device. IEEE Trans. Human-Machine Syst. 44, 731–742. 10.1109/THMS.2014.2348865.

20. Farajian, M., Leib, R., Kossowsky, H., Zaidenberg, T., Mussa-Ivaldi, F.A., Nisky, I., and Vaadia, E. (2020). Stretching the skin immediately enhances perceived stiffness and gradually enhances the predictive control of grip force. Elife 9. 10.7554/ELIFE.52653.

21. Thompson, S., Gregory, J.E., and Proske, U. (1990). Errors in force estimation can be explained by tendon organ desensitization. Exp. brain Res. 79, 365–372. 10.1007/BF00608246.

22. Luu, B.L., Day, B.L., Cole, J.D., and Fitzpatrick, R.C. (2011). The fusimotor and reafferent origin of the sense of force and weight. J. Physiol. 589, 3135. 10.1113/JPHYSIOL.2011.208447.

23. Roland, P.E., and Ladegaard-Pedersen, H. (1977). A quantitative analysis of sensations of tension and of kinaesthesia in man. Evidence for a peripherally originating muscular sense and for a sense of effort. Brain 100, 671–692. 10.1093/BRAIN/100.4.671.

24. McCloskey, D.I., Ebeling, P., and Goodwin, G.M. (1974). Estimation of weights and tensions and apparent involvement of a “sense of effort.” Exp. Neurol. 42, 220–232. 10.1016/0014-4886(74)90019-3.

25. Gandevia, S.C., Smith, J.L., Crawford, M., Proske, U., and Taylor, J.L. (2006). Motor commands contribute to human position sense. J. Physiol. 571, 703–710. 10.1113/jphysiol.2005.103093.

26. White, O., McIntyre, J., Augurelle, A.-S., and Thonnard, J.-L. (2005). Do novel gravitational environments alter the grip-force/load-force coupling at the fingertips? Exp. Brain Res. 163, 324–334. 10.1007/s00221-004-2175-8.

27. Descoins, M., Danion, F.R., and Bootsma, R.J. (2006). Predictive control of grip force when moving object with an elastic load applied on the arm. Exp. Brain Res. 172, 331–342. 10.1007/s00221-005-0340-3.

28. Augurelle, A.-S., Penta, M., White, O., and Thonnard, J.-L. (2003). The effects of a change in gravity on the dynamics of prehension. Exp. Brain Res. 148, 533–540. 10.1007/s00221-002-1322-3.

29. Opsomer, L., Théate, V., Lefèvre, P., and Thonnard, J.-L. (2018). Dexterous Manipulation During Rhythmic Arm Movements in Mars, Moon, and Micro-Gravity. Front. Physiol. 9, 1–10. 10.3389/fphys.2018.00938.

30. Wolpert, D.M., Ghahramani, Z., and Jordan, M.I. (1995). An internal model for sensorimotor integration. Science (80-.). 269, 1880–1882. 10.1126/science.7569931.

31. Takamuku, S., and Gomi, H. (2015). What you feel is what you see: Inverse dynamics estimation underlies the resistive sensation of a delayed cursor. Proc. R. Soc. B Biol. Sci. 10.1098/rspb.2015.0864.

32. Shadmehr, R., and Mussa-Ivaldi, F.A. (1994). Adaptive representation of dynamics during learning of a motor task. J. Neurosci. 14, 3208–3224. 10.1523/JNEUROSCI.14-05-03208.1994.

33. Flanagan, J.R., Nakano, E., Imamizu, H., Osu, R., Yoshioka, T., and Kawato, M. (1999). Composition and decomposition of internal models in motor learning under altered kinematic and dynamic environments. J. Neurosci. 19, RC34. citeulike-article-id:406855.

34. Bringoux, L., Macaluso, T., Sainton, P., Chomienne, L., Buloup, F., Mouchnino, L., Simoneau, M., and Blouin, J. (2020). Double-Step Paradigm in Microgravity: Preservation of Sensorimotor Flexibility in Altered Gravitational Force Field. Front. Physiol. 10.3389/fphys.2020.00377.

35. Casellato, C., Tagliabue, M., Pedrocchi, A., Papaxanthis, C., Ferrigno, G., and Pozzo, T. (2012). Reaching while standing in microgravity: A new postural solution to oversimplify movement control. Exp. Brain Res. 216, 203–215. 10.1007/s00221-011-2918-2.

36. Macaluso, T., Bourdin, C., Buloup, F., Mille, M.L., Sainton, P., Sarlegna, F.R., Vercher, J.L., and Bringoux, L. (2017). Sensorimotor reorganizations of arm kinematics and postural strategy for functional whole-body reaching movements in microgravity. Front. Physiol. 8, 1–12. 10.3389/fphys.2017.00821.

37. Papaxanthis, C., Pozzo, T., and McIntyre, J. (2005). Kinematic and dynamic processes for the control of pointing movements in humans revealed by short-term exposure to microgravity. Neuroscience 135, 371–383. 10.1016/j.neuroscience.2005.06.063.

38. Augurelle, A.-S. (2003). Feedback and feedforward processes underlying grip-load force coupling during cyclic vertical arm movements. UCLouvain.

39. Flanagan, J.R., and Wing, A.M. (1997). The role of internal models in motion planning and control: evidence from grip force adjustments during movements of hand-held loads. J. Neurosci. 17, 1519–1528. 10.1007/s00221-008-1691-3.

40. Nowak, D.A., Hermsdörfer, J., Schneider, E., and Glasauer, S. (2004). Moving objects in a rotating environment: rapid prediction of Coriolis and centrifugal force perturbations. Exp. Brain Res. 157, 241–254. 10.1007/s00221-004-1839-8.

41. Lafargue, G., Paillard, J., Lamarre, Y., and Sirigu, A. (2003). Production and perception of grip force without proprioception: Is there a sense of effort in deafferented subjects? Eur. J. Neurosci. 17, 2741–2749. 10.1046/j.1460-9568.2003.02700.x.

42. von Holst, E., and Mittelstaedt, H. (1950). The principle of reafference. Naturwissenschaften 3, 464–476.

43. Toffin, D., McIntyre, J., Droulez, J., Kemeny, A., and Berthoz, A. (2003). Perception and Reproduction of Force Direction in the Horizontal Plane. J. Neurophysiol. 90, 3040–3053. 10.1152/JN.00271.2003/ASSET/IMAGES/LARGE/9K1133482011.JPEG.

44. Van Beek, F.E., Tiest, W.M.B., and Kappers, A.M.L. (2013). Anisotropy in the haptic perception of force direction and magnitude. IEEE Trans. Haptics 6, 399–407. 10.1109/TOH.2013.37.

45. Delhaye, B.P., Schiltz, F., Barrea, A., Thonnard, J.-L., and Lefèvre, P. (2021). Measuring fingerpad deformation during active object manipulation. J. Neurophysiol. 126, 1455–1464. 10.1152/jn.00358.2021.

46. Sarlegna, F.R., Baud-Bovy, G., and Danion, F.R. (2010). Delayed Visual Feedback Affects Both Manual Tracking and Grip Force Control When Transporting a Handheld Object. J. Neurophysiol. 104, 641–653. 10.1152/jn.00174.2010.

47. Toma, S., Caputo, V., and Santello, M. (2020). Visual Feedback of Object Motion Direction Influences the Timing of Grip Force Modulation During Object Manipulation. Front. Hum. Neurosci. 14, 1–17. 10.3389/fnhum.2020.00198.

48. Grover, F.M., Schwab, S.M., and Riley, M.A. (2020). Grip Force-Load Force Coupling Is Influenced by Altered Visual Feedback about Object Kinematics. J. Mot. Behav. 52, 612–624. 10.1080/00222895.2019.1664977.

49. Gaveau, J., Berret, B., Angelaki, D.E., and Papaxanthis, C. (2016). Direction-dependent arm kinematics reveal optimal integration of gravity cues. Elife 5, 1–17. 10.7554/eLife.16394.

50. Crevecoeur, F., Thonnard, J.-L., and Lefèvre, P. (2009). Forward models of inertial loads in weightlessness. Neuroscience 161, 589–598. 10.1016/j.neuroscience.2009.03.025.

51. Mierau, A., Girgenrath, M., and Bock, O. (2008). Isometric force production during changed-Gz episodes of parabolic flight. Eur. J. Appl. Physiol. 102, 313–318. 10.1007/s00421-007-0591-8.

52. Sand, D.P., Girgenrath, M., Bock, O., and Pongratz, H. (2003). Production of isometric forces during sustained acceleration. Aviat. Sp. Environ. Med. 74, 633–637.

53. Girgenrath, M., Göbel, S., Bock, O., and Pongratz, H. (2005). Isometric force production in high Gz: Mechanical effects, proprioception, and central motor commands. Aviat. Sp. Environ. Med. 76, 339–343.

54. Lackner, J.R., and Graybiel, A. (1979). Parabolic flight: Loss of sense of orientation. Science (80-.). 206, 1105–1108. 10.1126/science.493998.

55. Clément, G.R., Moore, S.T., Raphan, T., and Cohen, B. (2001). Perception of tilt (somatogravic illusion) in response to sustained linear acceleration during space flight. Exp. Brain Res. 138, 410–418. 10.1007/s002210100706.

56. Young, L.R., Oman, C.M., Merfeld, D.M., Watt, D.G.D., Roy, S., DeLuca, C., Balkwill, D., Christie, J., Groleau, N., Jackson, D.K., et al. (1993). Spatial orientation and posture during and following weightlessness: Human experiments on spacelab life sciences. J. Vestib. Res. 3, 231–239.

57. Bock, O., Howard, I.P., Money, K.E., and Arnold, K.E. (1992). Accuracy of aimed arm movements in changed gravity. Aviat. Sp. Environ. Med. 63, 994–998.

58. Watt, D.G.D. (1997). Pointing at memorized targets during prolonged microgravity. Aviat. Sp. Environ. Med. 68, 99–103.

59. Fisk, J., Lackner, J.R., and DiZio, P. (1993). Gravitoinertial force level influences arm movement control. J. Neurophysiol. 69, 504–511. 10.1152/jn.1993.69.2.504.

60. Schneider, S., Brümmer, V., Göbel, S., Carnahan, H., Dubrowski, A., and Strüder, H.K. (2007). Parabolic flight experience is related to increased release of stress hormones. Eur. J. Appl. Physiol. 100, 301–308. 10.1007/s00421-007-0433-8.

61. Flanagan, J.R., and Beltzner, M.A. (2000). Independence of perceptual and sensorimotor predictions in the size-weight illusion. Nat. Neurosci. 10.1038/76701.

62. Grandy, M.S., and Westwood, D.A. (2006). Opposite perceptual and sensorimotor responses to a size-weight illusion. J. Neurophysiol. 10.1152/jn.00851.2005.

63. Buckingham, G., Cant, J.S., and Goodale, M.A. (2009). Living in A Material World: How Visual Cues to Material Properties Affect the Way That We Lift Objects and Perceive Their Weight. J. Neurophysiol. 102, 3111–3118. 10.1152/jn.00515.2009.

64. Lee, R., Klatzky, R.L., and Stetten, G.D. (2017). In-Situ Force Augmentation Improves Surface Contact and Force Control. IEEE Trans. Haptics 10, 545–554. 10.1109/TOH.2017.2696949.

65. Weber, B.M., and Stelzer, M. (2022). Sensorimotor impairments during spaceflight: Trigger mechanisms and haptic assistance. Front. Neuroergonomics 0, 24. 10.3389/FNRGO.2022.959894.

