## Supplementary_Figures for "A haptic illusion created by gravity"

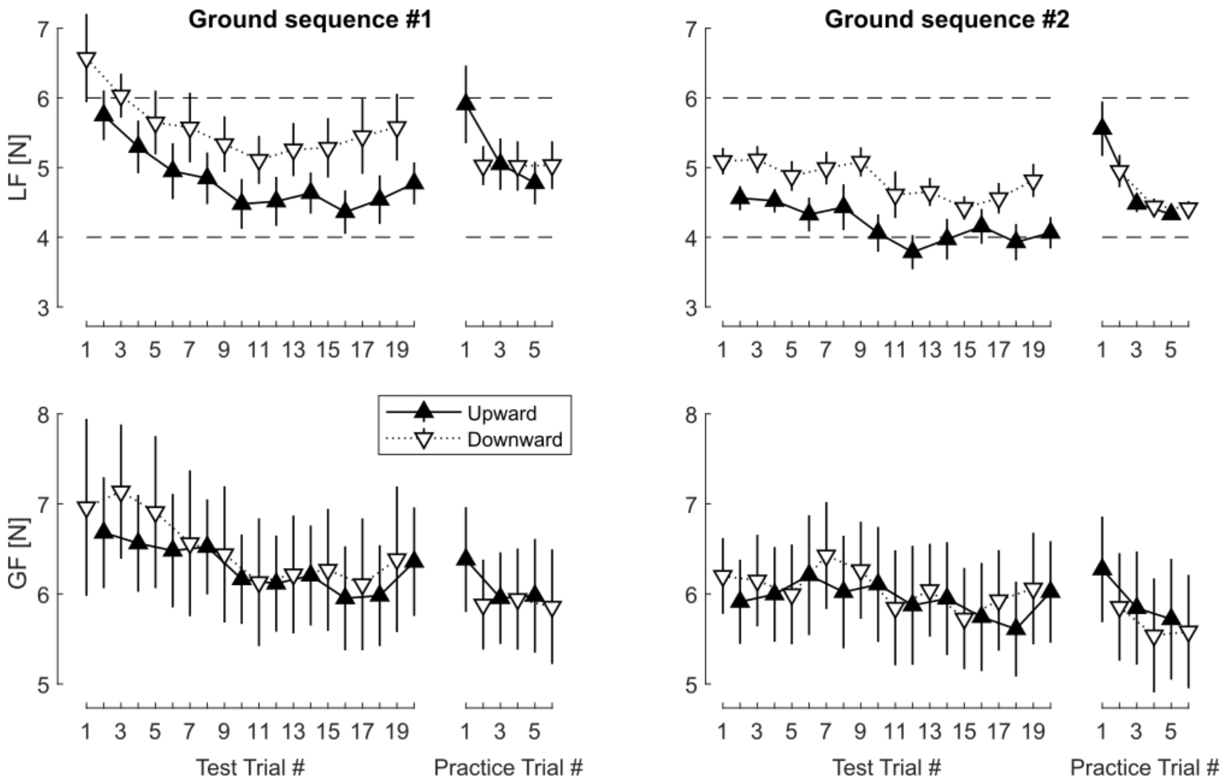


**Figure S1. Ground data**. Mean LF (upper panels) and GF (lower panels) values during the test and practice trials of the 1st (left) and 2nd (right) sequences performed on the ground. The dashed horizontal lines show the target boundaries for LF. Error bars show the mean ± SEM (N = 11).

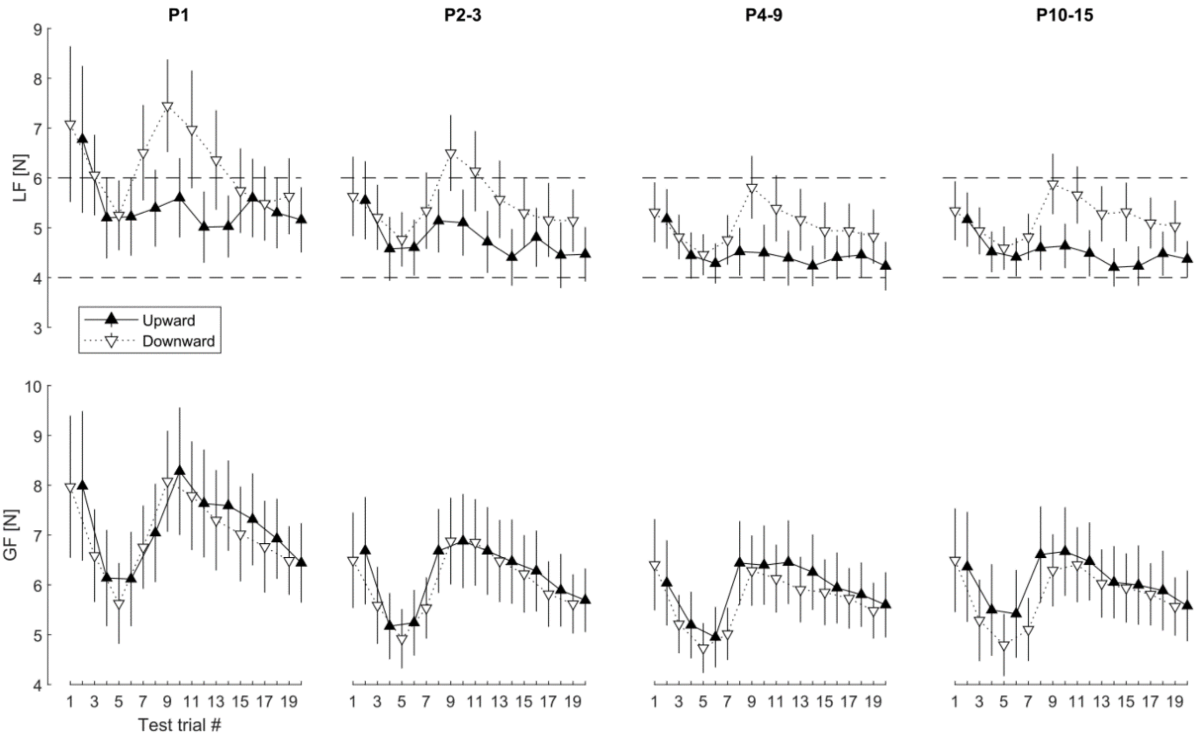


**Figure S2. Evolution of LF and GF across parabolas**. Mean LF and GF values obtained during test trials are shown for the first parabola (P1), parabolas 2–3, parabolas 4–9 and parabolas 10–15. Error bars show SEM (N = 11).

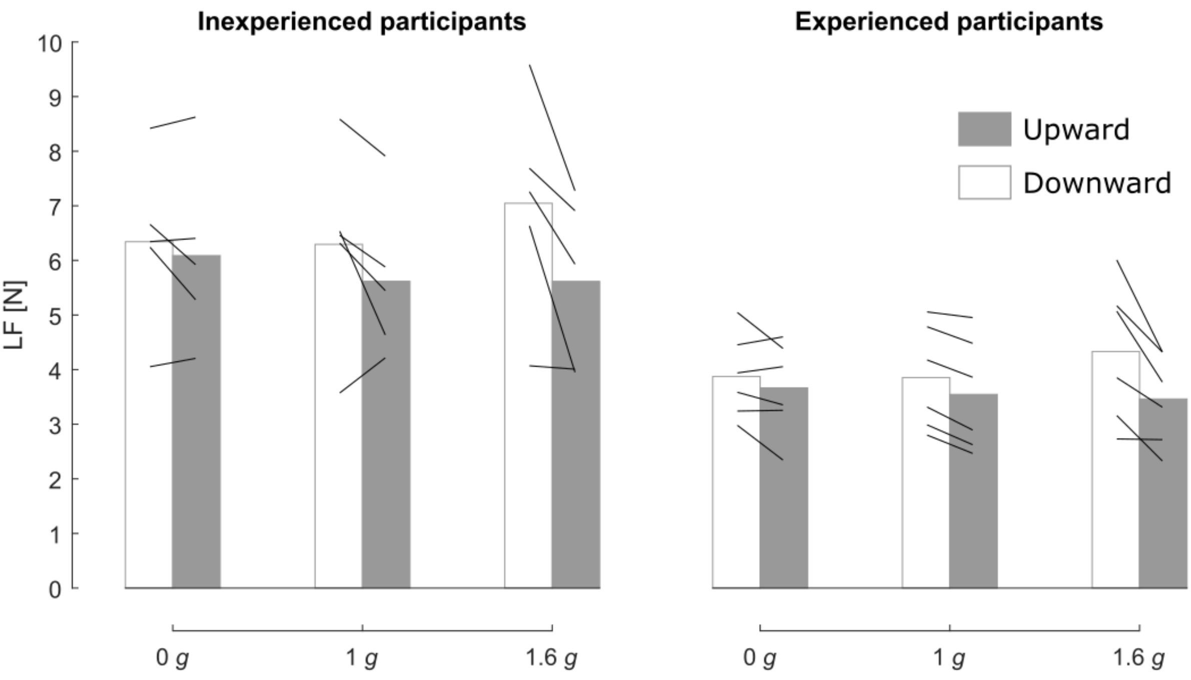


**Figure S3. Experienced versus inexperienced participants**. LF values obtained during the test trials for Inexperienced (N = 5) and Experienced (N = 6) participants (black lines). Inexperienced participants had never been exposed to parabolic flight before this study, whereas experienced ones had experienced at least one parabolic flight previously. The bars show the means across participants.
